# A single intranasal or intramuscular immunization with chimpanzee adenovirus vectored SARS-CoV-2 vaccine protects against pneumonia in hamsters

**DOI:** 10.1101/2020.12.02.408823

**Authors:** Traci L. Bricker, Tamarand L. Darling, Ahmed O. Hassan, Houda H. Harastani, Allison Soung, Xiaoping Jiang, Ya-Nan Dai, Haiyan Zhao, Lucas J. Adams, Michael J. Holtzman, Adam L. Bailey, James Brett Case, Daved H. Fremont, Robyn Klein, Michael S. Diamond, Adrianus C. M. Boon

**Affiliations:** Department of Internal Medicine, Washington University in Saint Louis School of Medicine, St. Louis, MO 63110, USA; Department of Molecular Microbiology and Microbial Pathogenesis, Washington University in Saint Louis School of Medicine, St. Louis, MO 63110, USA; Department of Pathology and Immunology, Washington University in Saint Louis School of Medicine, St. Louis, MO 63110, USA; Department of Biochemistry and Biophysics, Washington University in Saint Louis School of Medicine, St. Louis, MO 63110, USA; Department of Neuroscience, Washington University in Saint Louis School of Medicine, St. Louis, MO 63110, USA

## Abstract

The development of an effective vaccine against SARS-CoV-2, the etiologic agent of COVID-19, is a global priority. Here, we compared the protective capacity of intranasal and intramuscular delivery of a chimpanzee adenovirus-vectored vaccine encoding a pre-fusion stabilized spike protein (ChAd-SARS-CoV-2-S) in Golden Syrian hamsters. While immunization with ChAd-SARS-CoV-2-S induced robust spike protein specific antibodies capable or neutralizing the virus, antibody levels in serum were higher in hamsters immunized by an intranasal compared to intramuscular route. Accordingly, ChAd-SARS-CoV-2-S immunized hamsters were protected against a challenge with a high dose of SARS-CoV-2. After challenge, ChAd-SARS-CoV-2-S-immunized hamsters had less weight loss and showed reductions in viral RNA and infectious virus titer in both nasal swabs and lungs, and reduced pathology and inflammatory gene expression in the lungs, compared to ChAd-Control immunized hamsters. Intranasal immunization with ChAd-SARS-CoV-2-S provided superior protection against SARS-CoV-2 infection and inflammation in the upper respiratory tract. These findings support intranasal administration of the ChAd-SARS-CoV-2-S candidate vaccine to prevent SARS-CoV-2 infection, disease, and possibly transmission.

## INTRODUCTION

Severe acute respiratory syndrome coronavirus 2 (SARS-CoV-2) initiated a global pandemic in 2019, leading to millions of confirmed positive cases of coronavirus infection disease (COVID)-19, and an estimated case-fatality rate of 2-3% and infection fatality rate of 0.68% [1–3]. The elderly, immunocompromised and those with an underlying illness, including obesity, diabetes, hypertension, and chronic lung disease, are at greater risk of severe disease and death from SARS-CoV-2 [4, 5]. Antiviral therapies and vaccines are urgently needed to curb the spread of the virus and reduce infection and disease in the population.

The Golden Syrian hamster (*Mesocricetus auratus*) is one of several COVID-19 animal models [6–13]. Hamsters are naturally susceptible to SARS-CoV-2 infection, and intranasal inoculation results in mild-to-moderate disease including labored breathing, signs of respiratory distress, ruffled fur, weight loss and hunched posture [10, 13]. Aged and male hamsters develop more severe disease, mimicking COVID-19 in humans [11]. In hamsters, SARS-CoV-2 primarily infects the upper and lower respiratory tracts, although viral RNA and antigen has been detected in other tissues (e.g., intestines, heart, and olfactory bulb). The peak of virus replication occurs between days 2 and 3 post infection (dpi) and is cleared by 14 dpi in surviving animals. Histopathological analysis of infected hamsters shows multifocal interstitial pneumonia characterized by pulmonary consolidation starting as early as 2 dpi. Inflammation is associated with leukocyte infiltration, comprised primarily of macrophages and neutrophils, and an increase in type I and III interferon (IFN) and other pro-inflammatory cytokine and chemokines [14, 15]. High-resolution computed tomography scans shows airway dilation and consolidation in the lungs of infected hamsters [10]. SARS-CoV-2-induced lung pathology in hamsters appears driven by immune pathology, as lung injury is reduced in *STAT2^-/-^* hamsters despite an increase in viral burden and tissue dissemination [14].

The SARS-CoV-2 hamster model has been used to study the efficacy of several drugs and candidate vaccines. Hydroxychloroquine had no impact on infectious virus titers and disease, whereas favipiravir reduced viral burden only when high doses were used [16–19]. Several candidate vaccines also have been tested. Yellow fever 17D-vectored and adenovirus (Ad)26-vectored SARS-CoV-2 vaccine candidates conferred protection against SARS-CoV-2 challenge in hamsters [15, 20]. Hamsters immunized intramuscularly (IM) with Ad26-vectored prefusion-stabilized spike (S) protein sustained less weight loss and fewer SARS-CoV-2-infected cells in the lungs at 4 dpi [20]. Syrian hamsters immunized twice by intraperitoneal injection with YF17D-vector expressing the S protein of SARS-CoV-2 showed reduced viral burden, inflammatory gene expression, and pathology in the lung [15]. Other vectored vaccines including a Newcastle Disease virus-S (NDV-S) and vesicular stomatitis virus-S (VSV-S) vaccine delivered IM also protected Syrian hamsters from SARS-CoV-2 infection [21, 22]. Alternative routes of administration have not been tested in hamsters.

Here, we tested the efficacy of a chimpanzee adenovirus (ChAd)-vectored vaccine expressing a prefusion-stabilized version of the S protein of SARS-CoV-2 (ChAd-SARS-CoV-2-S, [23]) in Syrian hamsters following IM or intranasal (IN) delivery. A single dose of the vaccine induced a robust S protein specific antibody response capable of neutralizing SARS-CoV-2, with IN delivery inducing approximately 6-fold higher antibody titers than IM delivery. Upon challenge, the ChAd-SARS-CoV-2-S immunized animals had less infectious virus and viral RNA in the lungs and nasal swabs, and this was associated with reduced pathology and numbers of viral-infected cells in the lungs at 3 dpi. The upper respiratory tract, i.e. the nasal cavity, of the hamsters demonstrated reduced pathology and SARS-CoV-2 infected cells only after IN immunization with ChAd-SARS-CoV-2-S. Collectively, these data show differences in protection mediated by the same vaccine when alternative routes of immunization are used, and support intranasal vaccine delivery for optimal protection against SARS-CoV-2 challenge.

## MATERIALS AND METHODS

### SARS-CoV-2 infection of Golden Syrian hamsters

SARS-CoV-2 (strain 2019-nCoV/USA-WA1/2020) was propagated on MA-104 monkey kidney cells, and the virus titer was determined by focus forming and plaque assays. Five-week old male hamsters were obtained from Charles River Laboratories and housed at Washington University. Five days after arrival, a pre-immunization serum sample was obtained, and the animals (n = 10 per group) were vaccinated via intranasal (IN) or intramuscular (IM) route with 10^10^ viral particles of a chimpanzee adenovirus vector expressing a pre-fusion stabilized spike (S) protein of SARS-CoV-2 (ChAd-SARS-CoV-2-S [23]) or a control chimpanzee adenovirus vector (ChAd-Control) in 100 μL of phosphate buffered saline (PBS). Twenty-one days later, a second serum sample was obtained, and the animals were transferred to the enhanced biosafety level 3 laboratory. One day later, the animals were challenged via IN route with 2.5 × 10^5^ PFU of SARS-CoV-2. Animal weights were measured daily for the duration of the experiment. Two days after challenge, a nasal swab was obtained. The swab was moistened in 1.0 mL of serum-free media and used to rub the outside of the hamster nose. The swab was placed into the vial containing the remainder of the 1.0 mL of media, vortexed, and stored for subsequent virological analysis. Three days after challenge, a subset of animals was sacrificed, and their lungs were collected for virological and histological analysis. The left lobe was homogenized in 1.0 mL DMEM, clarified by centrifugation (21,000 × g for 5 minutes) and used for viral titer analysis by quantitative RT-PCR using primers and probes targeting the N gene or the 5’ UTR region, and by focus forming assay (FFA). From these same animals, we also collected serum for antibody analysis and heads for histological analysis. The remaining animals were sacrificed at 10 dpi, and serum was collected for analysis of antibody against the nucleoprotein (N protein) of SARS-CoV-2.

### Virus titration assays

FFA were performed on Vero-E6 cells in a 96-well plate. Lung tissue homogenates were serially diluted 10-fold, starting at 1:10, in cell infection medium (DMEM + 2% FBS + L-glutamine + penicillin + streptomycin), and 100 μl of the diluted virus was added to two wells per dilution per sample. After 1 h at 37°C, the inoculum was aspirated, the cells were washed with PBS, and a 1% methylcellulose overlay in infection medium was added. Positive and negative controls were included in every assay. Twenty-four hours after virus inoculation, the cells were fixed with formalin, and infected cells were detected by the addition of 100 μL of 1:1000 diluted anti-S protein monoclonal antibody (1C02, gift from Dr. Ellebedy at Washington University) in permeabilization buffer (1x PBS, 2% FBS, 0.2% saponin (Sigma, Cat #S7900)) for 1 h at 20°C or overnight at 4°C, followed by an anti-human-IgG-HRP antibody (Sigma, Cat. #A6029) in permeabilization buffer for 1 h at 20°C. The assay was developed using TMB substrate (Vector laboratories, SK4400) for 5-10 min at 20°C. The assay was stopped by washing the cells with water. The number of foci per well were counted on the BioSpot analyzer (Cellular Technology Limited) and used to calculate the focus forming units/mL (FFU/mL).

Plaque assays were performed on Vero E6 cells in 24-well plates. Nasal swabs or lung tissue homogenates were serially diluted 10-fold, starting at 1:10, in cell infection medium (DMEM + 2% FBS + L-glutamine + penicillin + streptomycin). Two hundred and fifty microliters of the diluted virus were added to a single well per dilution per sample. After 1 h at 37°C, the inoculum was aspirated, the cells were washed with PBS, and a 1% methylcellulose overlay in MEM supplemented with 2% FBS was added. Seventy-two hours after virus inoculation, the cells were fixed with 4% formalin, and the monolayer was stained with crystal violet (0.5% w/v in 25% methanol in water) for 1 h at 20°C. The number of plaques were counted and used to calculate the plaque forming units/mL (PFU/mL).

To quantify viral load in nasal swabs and lung tissue homogenates, RNA was extracted using RNA isolation kit (Omega). SARS-CoV-2 RNA levels were measured by one-step quantitative reverse transcriptase PCR (qRT-PCR) TaqMan assay as described previously [23]. A SARS-CoV-2 nucleocapsid (N) specific primers/probe set (L primer: ATGCTGCAATCGTGCTACAA; R primer: GACTGCCGCCTCTGCTC; probe: 5’-FAM/TCAAGGAAC/ZEN/AACATTGCCAA/3’-IABkFQ) or 5’ UTR specific primers/probe set (L primer: ACTGTCGTTGACAGGACACG; R primer: AACACGGACGAAACCGTAAG; probe: 5’-FAM/CGTCTATCT/ZEN/TCTGCAGGCTG/3’-IABkFQ). Viral RNA was expressed as (N) gene or 5’ UTR copy numbers per mg for lung tissue homogenates or mL for nasal swabs, based on a standard included in the assay, which was created via *in vitro* transcription of a synthetic DNA molecule containing the target region of the N gene and 5’-UTR region.

### ELISA

Purified viral antigens (S, RBD, or NP) [23] were coated onto 96-well Maxisorp clear plates at 2 μg/mL in 50 mM Na_2_CO_3_ pH 9.6 (70 μL) or PBS (50 μL) overnight at 4°C. Coating buffers were aspirated, and wells were blocked with 200 μL of 1X PBS + 0.05% Tween-20 + 5% BSA + 0.02% NaN_3_ (Blocking buffer, PBSTBA) or 1X PBS + 0.05% Tween-20 + 10% FCS (PBSTF) either for 2 h at 20°C, 1 h at 37°C or overnight at 4°C. Heat-inactivated serum samples were diluted in PBSTBA or PBSTF in a separate 96-well polypropylene plate. The plates then were washed thrice with 1X PBS + 0.05% Tween-20 (PBST), followed by addition of 50 μL of respective serum dilutions. Sera were incubated in the blocked ELISA plates for at least 1 h at room temperature. The ELISA plates were again washed thrice in PBST, followed by addition of 50 μL of 1:1000 anti-hamster-IgG(H+L)-HRP (Southern Biotech Cat. #6061-05) in PBST or PBSTF or 1:1000 anti-hamster-IgG2/IgG3-HRP in PBST or PBSTF (Southern Biotech Cat. #1935-05). Plates were incubated at room temperature for 1-2 h, washed thrice in PBST and 50 μL of 1-Step Ultra TMB-ELISA was added (Thermo Fisher Scientific, Cat. #34028). Following a 12 to 15-min incubation, reactions were stopped with 50 μL of 2 M H_2_SO_4_. The absorbance of each well at 450 nm was read (Synergy H1 or Epoch) within 2 min of addition of H_2_SO_4_.

### SARS-CoV-2 neutralization assay

Heat-inactivated serum samples were diluted 1:10 fold serially and incubated with 10^2^ FFU of SARS-CoV-2 for 1 h at 37°C. The virus-serum mixtures were added to Vero-E6 cell monolayers in 96-well plates and incubated for 1 h at 37°C. Subsequently, cells were overlaid with 1% (w/v) methylcellulose in MEM supplemented with 2% FBS. Plates were incubated for 30 h before fixation using 4% PFA in PBS for 1 h at 20°C. Cells were washed and then sequentially incubated with anti-SARS-CoV-2 CR3022 mAb [24] (1 μg/mL) and a HRP-conjugated goat anti-human IgG (Sigma, Cat#A6029) in PBS supplemented with 0.1% (w/v) saponin and 0.1% BSA. TrueBlue peroxidase substrate (KPL) was used to develop the plates before counting the foci on a BioSpot analyzer (Cellular Technology Limited).

### Histology and RNA *in situ* hybridization

The lungs and heads from SARS-CoV-2 infected and control hamsters were fixed in 10% formalin for seven days. Lungs were embedded in paraffin and sectioned before hematoxylin and eosin staining (H & E) and RNA *in situ* hybridization (RNA-ISH) to detect SARS-CoV-2 RNA. Following formalin fixation, heads were decalcified in 0.5 M EDTA for seven days, cryoprotected in three exchanges of 30% sucrose for three days, and then embedded in O.C.T. compound before RNA-ISH. RNA-ISH was performed using a probe against the S gene of SARS-CoV-2 (V-nCoV2019-S, Cat #848561) with the RNAscope® 2.5 HD Assay— BROWN (ACDBio, Cat#322310) according to the manufacturers’ recommendations. Lung slides were scanned using the Hamamatsu NanoZoomer slide scanning system and head sections were imaged using the Zeiss AxioImager Z2 system. Lung sections were scored according to a previous publication [10] (<10% affected lung tissue = 1, >10% but <50% affected area = 2, >50% affected area = 3).

### Host response gene analysis

RNA extracted from hamster lung tissue homogenates was used to synthesize cDNA using random hexamers and Superscript III (Thermo Scientific) with the addition of RNase inhibitor according to the manufacturer’s protocol. The expression of 22 inflammatory host genes was determined using PrimeTime Gene Expression Master Mix (Integrated DNA Technologies) with primers/probe sets specific for *C3, C5, Ccl2, Ccl3, Ccl5, Csf3, Cxcl10, Ddx58, Ifit3, Ifng, Irf7, IL1b, IL4, IL5, IL6, IL7, IL10, IL12p40, IL15, Stat1, Stat6*, and *Tnfa* and results were normalized to *Rpl18* and *B2m* levels. The primers and probes were derived from previous publications [25] or developed in-house (see **Table S1**). Fold change was determined using the 2-ΔΔCt method comparing immunized and SARS-CoV-2 challenged hamsters to naïve controls.

### Statistical Analysis

The data was analyzed with GraphPad Prism 9.0 and statistical significance was assigned when *P* values were < 0.05. All tests and values are indicated in the relevant Figure legends.

## RESULTS

### Development of the SARS-CoV-2 hamster model

To establish the utility of the hamster model in our hands, we inoculated twenty-one 5-6 week old male hamsters IN with 2 × 10^5^ plaque forming units (PFU) of a fully infectious SARS-CoV-2 isolate in 100 μL of PBS. A control group of three 5-6 week old male hamsters was inoculated with PBS. Mock-infected animals continued to gain weight at a rate of ~2.4 grams or ~3% per day (**Fig 1A**). In contrast, SARS-CoV-2 inoculated hamsters began to lose weight at 2 dpi, and this continued through days 4-5, at which point the animals had lost approximately 10% of their body weight (**Fig 1A**). This decrease was associated with a reduction in food intake between 1 and 4 dpi (**Fig 1B**).

**Figure 1:**
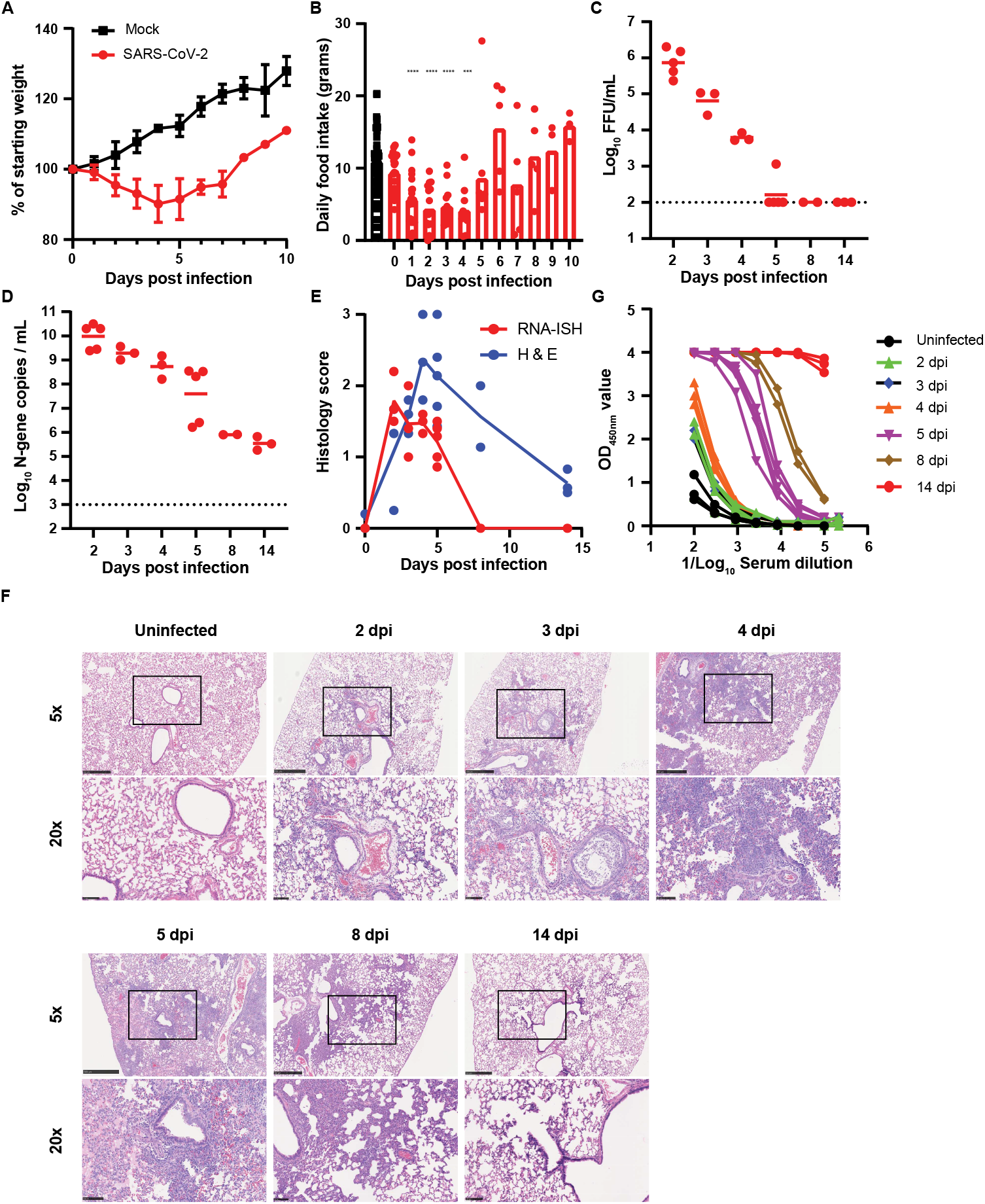
Development of the SARS-CoV-2 hamster model. (**A**) Mean + standard deviation (SD) weight loss or weight gain of uninfected (n = 3) or SARS-CoV-2 infected (n = 21). (**B**) Daily food intake of uninfected and infected hamsters. Data points for the uninfected hamsters (n = 3 per day) were recorded for 14 days and plotted. For infected hamsters, food intake 1 to 10 dpi was recorded (**** *P* < 0.0001, *** *P* < 0.001 by ANOVA with Dunnett’s multiple comparison against the uninfected hamsters). (**C**) Infectious virus titer was quantified by FFA from homogenates of the left lung lobe at indicated time points. Each dot is an individual hamster and bars indicate median values (dotted line is the limit of detection). (**D**) Lung viral RNA was quantified in the left lung lobe at indicated time points after infection. Each dot is an individual hamster and bars indicate median values (dotted line is the limit of detection). (**E**) Pathology score of the lungs from infected hamsters. <10% affected = 1, >10% but <50% = 2, >50% = 3. Each lobe was scored, and the average score was plotted per animals. The solid line is the average score per day for RNA *in situ* hybridization (red line and dots) or inflammation (blue line and dots). (**F**) Representative images at 5x and 20x magnification of H & E staining of SARS-CoV-2 infected hamsters sacrificed at different time points after inoculation (n = 5 for 2 dpi, n = 3 for 3 dpi, n = 3 for 4 dpi, n = 5 for 5 dpi, n = 2 for 8 dpi, n = 3 for 14 dpi, n = 3 for uninfected). (**G**) Serum S protein specific IgG(H+L) responses in SARS-CoV-2 infected hamsters. Each color is a different day after infection. (C-D) Bars indicate median values, and dotted lines are the LOD of the assays.

At indicated time points after infection, hamsters were sacrificed, and tissues were collected for analysis of viral burden, histology, and serological response. The left lung lobe was collected, homogenized, and used to quantify SARS-CoV-2 N-gene copy number and infectious virus titer by qPCR and focus-forming assay (FFA), respectively. Infectious virus titers peaked at 2 dpi with 8 × 10^5^ focus forming units/mL (FFU/mL), and levels declined to low or undetectable by 5 dpi (**Fig 1C**). SARS-CoV-2 N-gene copy number also peaked at 2 dpi at 10^10^ copies per μL and gradually declined to 10^5^-10^6^ copies by 8 to 14 dpi (**Fig 1D**). The remainder of the lung tissue was fixed in formalin, embedded, and sectioned for viral RNA *in situ* hybridization (ISH) and hematoxylin and eosin (H & E) staining. SARS-CoV-2 viral RNA was detected by RNA-ISH at 2-5 dpi (**Fig 1E** and **Fig S1**) and was no longer detectable by 8 dpi. Viral RNA was localized to both airway and alveolar epithelial cells (**Fig S1**). Infection was accompanied by immune cell infiltration in peribronchiolar and adjacent alveolar locations from 2 through 8 dpi (**Fig 1E-F**), a pattern that is consistent with bronchopneumonia. The immune cell infiltration was associated with alveolar edema, exudate, tissue damage and intraparenchymal hemorrhage (**Fig 1E-F**). Each of these features of histopathology were markedly decreased by 14 dpi (**Fig 1F**).

Serum samples were assayed for the presence of antibodies specific for purified, recombinant S protein by ELISA. Low or undetectable antibody responses were detected through 4 dpi (**Fig 1G**. By day 5, S-specific IgG(H+L) responses were detected in all five animals, and the serum antibody titer further increased between 8 and 14 dpi.

### Chimpanzee Ad-vectored vaccine elicits robust antibody responses against SARS-CoV-2 in hamsters

We assessed the immunogenicity of a replication-incompetent ChAd vector encoding a prefusion-stabilized, full-length sequence of SARS-CoV-2 S protein (ChAd-SARS-CoV-2-S) [23] in Golden Syrian hamsters. We used a ChAd vector without a transgene (ChAd-control) as a control. Groups of ten 5-6 week-old male hamsters were immunized once via IN or IM route with 10^10^ virus particles of ChAd-control or ChAd-SARS-CoV-2-S. Serum was collected prior to immunization or 21 days after, and antibody responses were evaluated by ELISA against purified recombinant S and RBD proteins. Immunization with ChAd-SARS-CoV-2-S induced high levels of anti-S and anti-RBD IgG(H+L) and IgG2/IgG3 antibodies 21 days later, whereas low or undetectable levels of S- and RBD-specific antibodies were present in samples from ChAd-control immunized animals (**Fig 2A-F** and **Fig S2**). The antibody response was significantly higher after IN than IM immunization (5 to 7-fold, *P* < 0.0001 for anti-S and anti-RBD respectively, **Fig 2G-H**). Serum samples also were tested for neutralization of infectious SARS-CoV-2 by focus-reduction neutralization test (FRNT). As expected, pre-immunization sera or sera from hamsters immunized with ChAd-control did not inhibit virus infection (**Fig 2I**). In contrast, sera from animals immunized with ChAd-SARS-CoV-2-S neutralized infectious virus with geometric mean titers (GMT) of 1:1217 and 1:276 for IN and IM immunization routes, respectively (**Fig 2I**).

**Figure 2:**
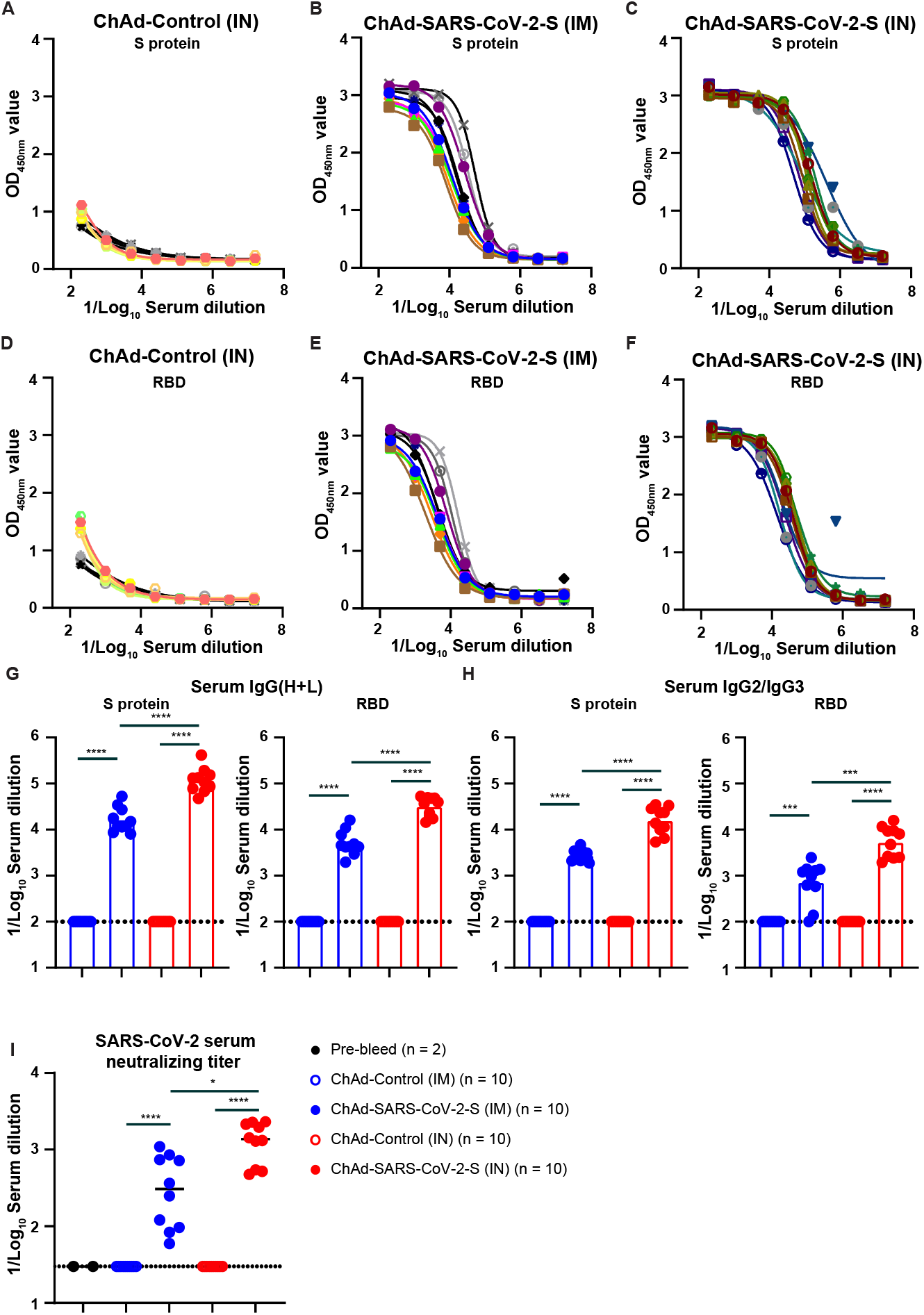
Humoral immune response following IN and IM immunization. (**A-C**) Anti-S protein-specific serum IgG(H+L) titers in hamsters immunized IN with ChAd-Control (**A**) or with ChAd-SARS-CoV-2-S IM (**B**) or IN (**C**). Each line is an individual animal. (**D-G**) Receptor binding domain (RBD)-specific serum IgM titers in hamsters immunized IN with ChAd-Control (**D**), or with ChAd-SARS-CoV-2-S IM (**E**) or IN (**F**). Each line is an individual animal. (**G-H**) IC_50_ values for S protein specific or RBD specific IgG(H+L) (**G**) or IgG2/IgG3 (**H**) serum antibodies in hamsters vaccinated IM (blue symbols) or IN (red symbols) with ChAd-Control (open symbols) or ChAd-SARS-CoV-2-S (closed symbols). (**** *P* < 0.0001, *** *P* < 0.001 by Mann-Whitney test with a Bonferroni correction for multiple comparisons). (**I**) SARS-CoV-2 serum neutralizing titer, measured by FRNT, in hamsters vaccinated IM or IN with ChAd-Control or ChAd-SARS-CoV-2-S. (**** *P* < 0.0001, * *P* < 0.05 by Mann-Whitney test with a Bonferroni correction for multiple comparisons). (**G-I**) Bars indicate median values, and dotted lines are the LOD of the assays.

### Immunization with Chimpanzee Ad-vectored vaccine protects hamsters from SARS-CoV-2 challenge

We next evaluated the protective effect of the ChAd vaccines in the hamster SARS-CoV-2 challenge model. Golden Syrian hamsters immunized with ChAd-SARS-CoV-2-S or ChAd-control were challenged IN with 2.5 × 10^5^ PFU of SARS-CoV-2, and 2 dpi a nasal swab was collected for viral burden analysis by qPCR and plaque assay. The N-gene copy number in the ChAd-control immunized animals was ~10^9^ copies per mL in both the IM and IN control groups (**Fig 3A**). Immunization with ChAd-SARS-CoV-2-S reduced the N-gene copy number by 100-fold in the IN (10^7^/mL) and 10-fold in the IM (10^8^/mL) immunized animals (*P* < 0.0001 and *P* < 0.001 respectively, **Fig 3A**). N-gene copy number was significantly lower in the IN than IM immunized animals (*P* < 0.05, **Fig 3A**). At 2 dpi, infectious virus was detected by plaque assay in 4 of 20 ChAd-SARS-CoV-2-S-immunized animals and 15 of 20 ChAd-control immunized animals (**Fig 3B**). At 3 dpi, six hamsters per group were sacrificed, and lungs were collected for viral burden analysis (left lobe) by qPCR or FFA, or for histology (other lung lobes). In the control groups, we detected 10^9^-10^10^ copies of N-gene per mg of lung homogenate, and the mean infectious titer was 6 × 10^4^ FFU/mL. No difference was observed between the two control groups. Immunization with ChAd-SARS-CoV-2-S vaccine significantly reduced the N-gene copy number (*P* < 0.01, **Fig 3C**) and infectious titer in both the IN and IM immunized group (**Fig 3D**). A comparison between IN and IM immunization revealed significantly lower N-gene copies per mg (788-fold, *P* < 0.01, **Fig 3C**), but not in infectious virus titer (*P* = 0.5, **Fig 3D**), in the lungs of IN immunized animals. The remaining four animals per group were monitored for weight loss for 10 days. The ChAd-control immunized animals lost an average of 4% and 8% of their starting body weight (**Fig 3E**). Immunization with ChAd-SARS-CoV-2-S attenuated weight loss after SARS-CoV-2 challenge in both groups (*P* < 0.01, **Fig 3E**), with a possibly greater effect following IN immunization. To assess if the ChAd-SARS-CoV-2-S vaccine induced sterilizing immunity by either of the immunization routes, we collected the serum 10 dpi to test for the presence of antibodies against recombinant NP by ELISA. A robust anti-NP IgG(H+L) (**Fig S3**) and IgG2/3 (**Fig 3F**) antibody response was detected in all ChAd-control and ChAd-SARS-CoV-2-S immunized animals.

**Figure 3:**
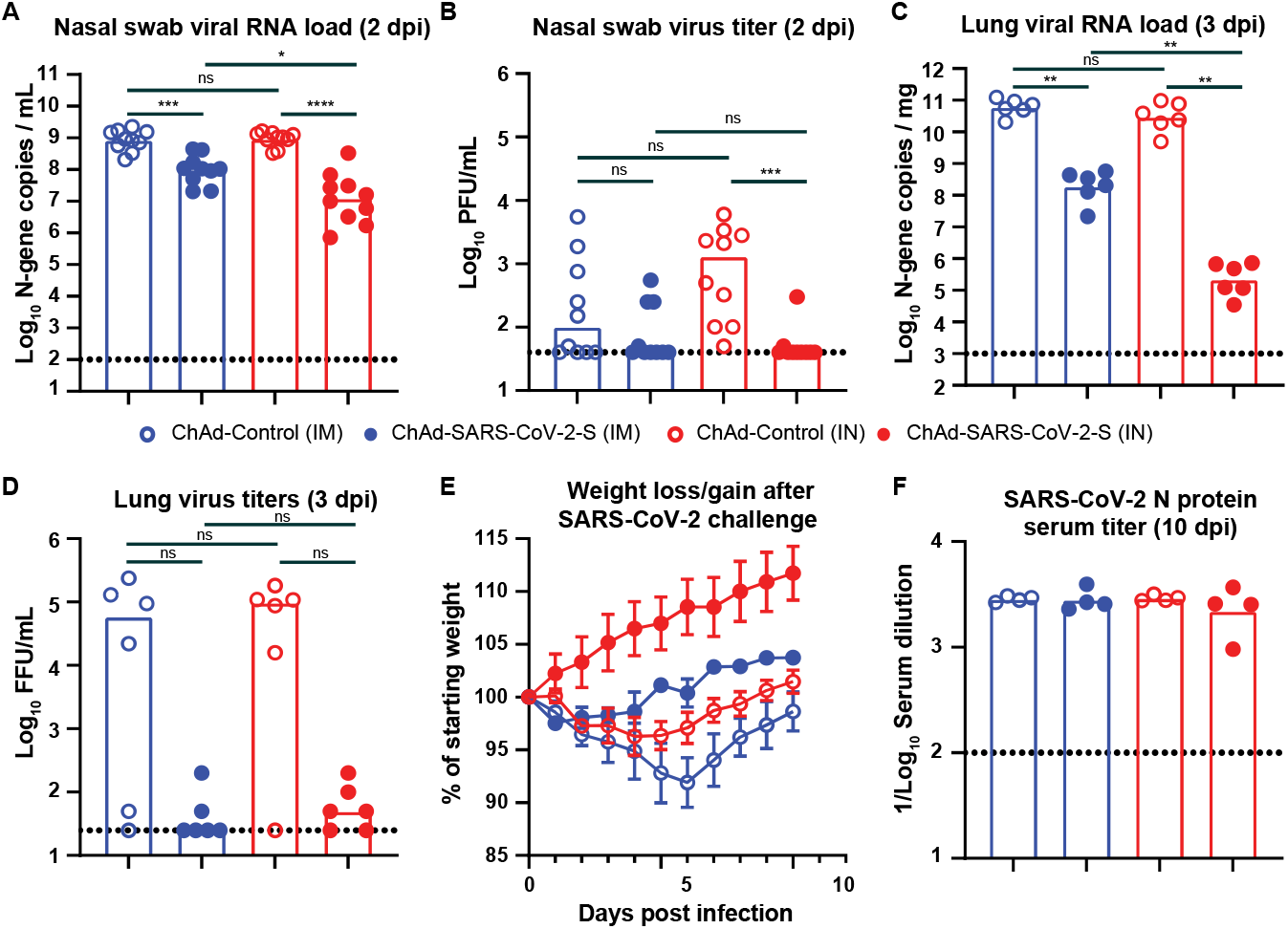
IN immunization offers superior protection against challenge with SARS-CoV-2. Twenty-eight days after a single IM (blue symbols) or IN (red symbols) vaccination with ChAd-Control (open symbols) or ChAd-SARS-CoV-2-S (closed symbols), hamsters were challenged with 2.5 × 10^5^ PFU of SARS-CoV-2, and nasal swabs (**A** and **B**) and lungs (**C** and **D**) were collected for analysis of viral RNA levels by qPCR (**A** and **C**) and infectious virus by plaque assay (**B** and **D**). (**** *P* < 0.0001, *** *P* < 0.001, ** *P* < 0.01, * *P* < 0.05, ns = not significant by Mann-Whitney test with a Bonferroni correction for multiple comparisons). (**E**) Mean + SD of weight loss/gain in SARS-CoV-2 challenged hamsters (n = 4 per group). (**F**) SARS-CoV-2 N protein serum titer, measured by ELISA, in hamsters vaccinated IM or IN with ChAd-Control or ChAd-SARS-CoV-2-S. (**A-D** and **F**) Bars indicate median values, and dotted lines are the limit of detection of the assays.

### Immunization with Chimpanzee Ad-vectored vaccine minimizes lung pathology in hamsters

To support these findings, we performed RNA *in situ* hybridization (RNA-ISH) and H & E staining on sections from formalin-fixed lung tissues from immunized hamsters. RNA-ISH detected viral RNA in all animals immunized with ChAd-control vaccine (**Fig 4A-B** and **Fig S4**). On average, ~20% of the section was positive for SARS-CoV-2 RNA. The presence of viral RNA was associated with inflammation, tissue damage, and bronchopneumonia, as evidenced by immune cell infiltration around bronchioles, alveolar edema, fluid exudates, and intraparenchymal hemorrhage (**Fig S4**). In contrast, sections from animals immunized IM with ChAd-SARS-CoV-2-S contained no or few SARS-CoV-2 positive cells by RNA-ISH and inflammation was greatly reduced (*P* < 0.01, **Fig 4A-B** and **Fig S5**). No SARS-CoV-2 positive cells were detected following IN immunization with ChAd-SARS-CoV-2-S (*P* < 0.01, **Fig 4A-B** and **Fig S5**).

**Figure 4:**
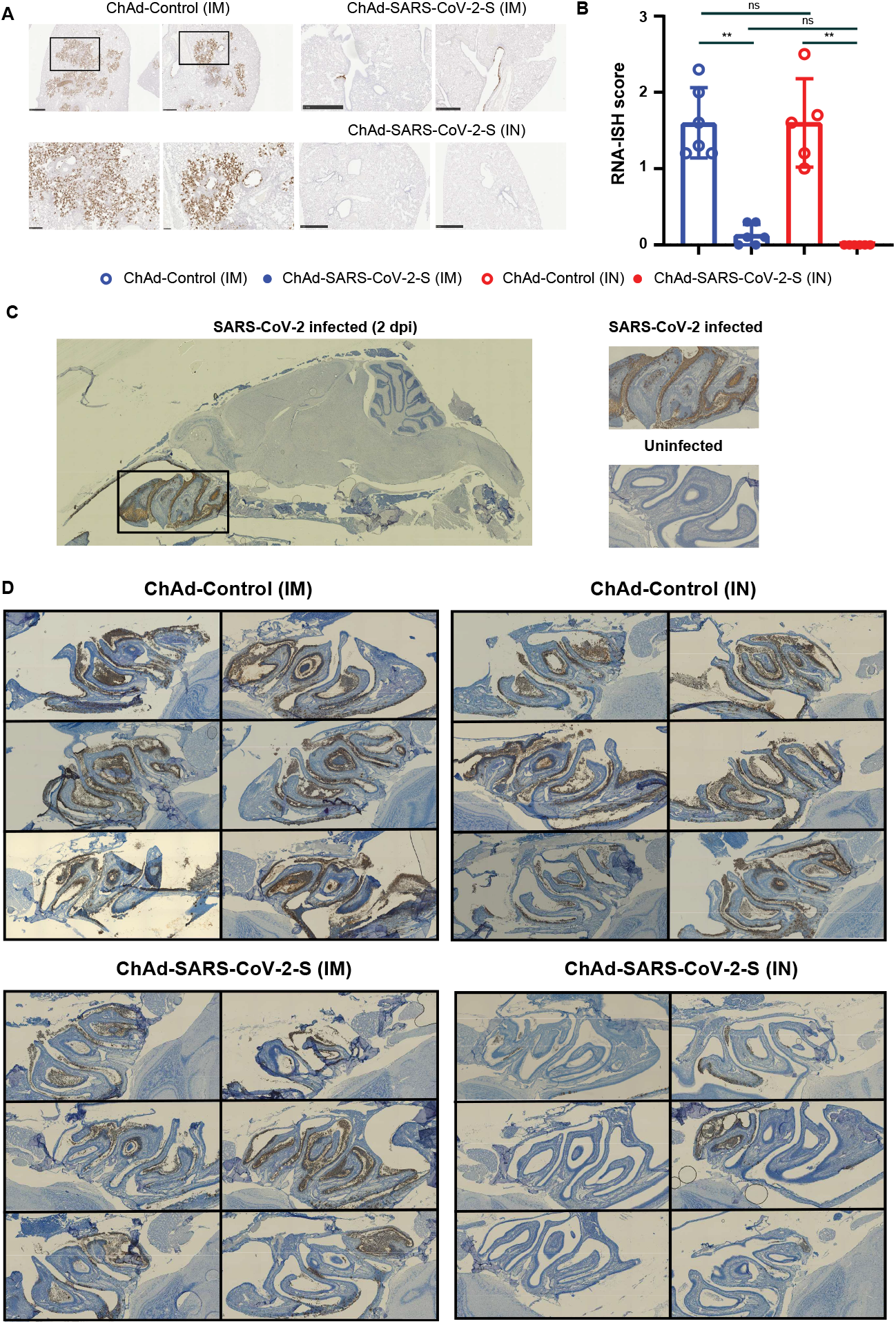
IN immunization offers superior protection against SARS-CoV-2 infection of the nasal epithelium. (**A**) RNA *in situ* hybridization (ISH) for SARS-CoV-2 viral RNA in hamster lung sections. Representative images of the ChAd-Control (IM), ChAd-SARS-CoV-2-S (IM) and ChAd-SARS-CoV-2-S (IN) sections. (**B**) Comparison of RNA-ISH staining between groups of hamsters. Each lobe was scored according to the following system, <10% RNA-positive = 1, >10% but <50% RNA-positive = 2, >50% RNA-positive = 3, and the average score was plotted per animals. (** *P* < 0.01, ns = not significant by Mann-Whitney U test with a Bonferroni correction for multiple comparisons). (**C**) Representative images of sagittal sections of hamster heads infected with SARS-CoV-2 for 2 days or uninfected control. RNA-ISH was performed on the sections and SARS-CoV-2 viral RNA was detected in the nasal turbinate. (**D**) Detection of SARS-CoV-2 viral by RNA-ISH in sagittal sections of hamster heads from the immunized and SARS-CoV-2 challenged animals.

An ideal SARS-CoV-2 vaccine would confer protection against disease and prevent virus infection and transmission. We hypothesized that IN delivery of the vaccine could provide superior protection in the upper respiratory tract compared to IM delivery. Hamster heads were collected 3 dpi and fixed in formalin. Following decalcification and embedding, sagittal sections were obtained and RNA-ISH was performed (**Fig 4C**). SARS-CoV-2 RNA was detected in the nasal cavity and ethmoturbinates of all 12 hamsters immunized IN or IM with ChAd-control (**Fig 4D, left panel**). No difference in viral RNA staining was observed between the two groups. Animals immunized IM with ChAd-SARS-CoV-2-S contained fewer SARS-CoV-2 positive cells and less cellular debris than ChAd-Control vaccinated animals (**Fig 4D**). However, animals immunized IN with ChAd-SARS-CoV-2-S had the fewest number of SARS-CoV-2-positive cells, and cellular debris was further reduced (**Fig 4D**). Collectively, these studies show that the ChAd-SARS-CoV-2-S vaccine is highly protective in the hamster model of COVID-19, and IN delivery of this vaccine provides superior protection against upper respiratory tract infection.

### Inflammatory gene expression is reduced after SARS-CoV-2 challenge in the ChAd-SARS-CoV-2 immunized hamsters

Lung pathology after SARS-CoV-2 infection appears to be driven by inflammation [14]. Thus, a successful vaccine should reduce or eliminate the inflammatory response after infection or challenge with SARS-CoV-2. The inflammatory response was evaluated in the ChAd-control and ChAd-SARS-CoV-2-S immunized hamsters 3 days after challenge with SARS-CoV-2. RNA was extracted from the tissue homogenates and analyzed by qRT-PCR using 24 different primer-probe sets specific for two housekeeping genes (*ß2m* and *Rpl18*) and 22 different innate and inflammatory host genes (**Table S1**). Compared to five naïve animals, expression of 8/22 inflammatory host genes (*Ccl2, Ccl3, Cxcl10, Ddx58, Ifit3, IL10, IL12p40*, and *Irf7* increased > 2-fold in the ChAd-Control immunized and SARS-CoV-2 challenged animals (**Fig 5A**). A significant increase in gene-expression was observed for *Ccl3, Ifit3, Cxcl10*, and *Irf7* in both ChAd-Control-IN and ChAd-Control-IM animals (*P* < 0.05, **Fig 5A**). ChAd-SARS-CoV-2-S immunization significantly reduced inflammatory gene expression after SARS-CoV-2 challenge (**Fig 5B**) with the expression of a subset of host genes, such as *Ccl5, Ccl3* and *Cxcl10*, near normal levels. A comparison in host gene expression between IM and IN immunization identified *Irf7* and *Ifit3* as two host genes whose expression was significantly (*P* < 0.01) lower in the IN compared to IM immunized animals (**Fig 5B**). These data suggest that IN delivery of the ChAd-vectored SARS-CoV-2-S vaccine provides greater protection against SARS-CoV-2 infection, inflammation, and disease in hamsters.

**Figure 5:**
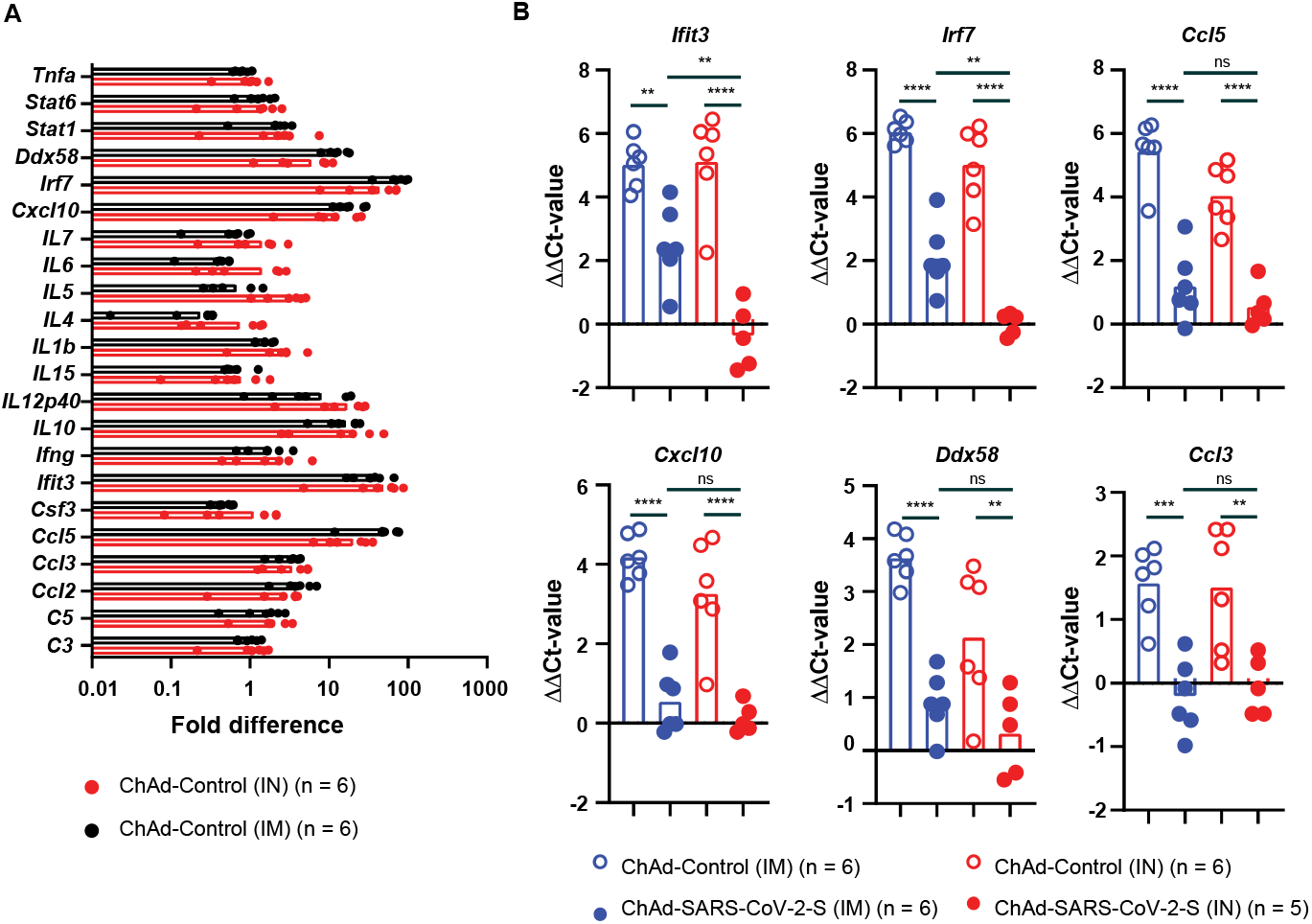
ChAd-SARS-CoV-2 immunization ameliorates inflammatory gene expression following SARS-CoV-2 challenge. Inflammatory gene-expression (n = 22) was quantified by RT-PCR in RNA extracted from lung homogenates 3 dpi (primer and probe sequences are in Table 1). (**A**) Fold increase in gene-expression for ChAd-Control immunized (IN in red and IM in black) and SARS-CoV-2 challenged hamsters. (**B**) ΔΔCt-values for *Ifit3, Irf7, Ccl5, Cxcl10, Ddx58* and *Ccl3* in ChAd-Control (open symbols) and ChAd-SARS-CoV-2-S (closed symbols) immunized and SARS-CoV-2 challenged animals 3 dpi. (ns = not significant, **** *P* < 0.0001, *** *P* < 0.001, ** *P* < 0.01 by one-way ANOVA with a Šidák correction for multiple comparisons). Each dot is an individual animal from two experiments. Bars indicate average values.

## DISCUSSION

Effective vaccines against SARS-CoV-2 are needed to combat the devastating pandemic. In this study, we evaluated IN and IM delivery of a ChAd-vectored vaccine expressing a prefusion stabilized S protein of SARS-CoV-2 in the Syrian hamster challenge model. A single dose of ChAd-SARS-CoV-2 induced S- and RBD-specific serum antibodies capable of neutralizing SARS-CoV-2. Antibody responses were higher after IN than IM immunization. Following challenge with a high dose of SARS-CoV-2, IN and IM immunization reduced infectious virus titers and viral RNA levels in the lungs and nasal swabs, albeit the effect was greater following IN immunization. Immunization with ChAd-SARS-CoV-2 also reduced weight loss, lung pathology and inflammatory gene expression in the lungs of the animals with a greater effect again seen after IN immunization. Finally, IN immunization protected the upper respiratory tract of hamsters, whereas IM immunization did not. Combined, these studies demonstrate that a single dose of ChAd-SARS-CoV-2-S vaccine delivered IN provides better protection than IM immunization against SARS-CoV-2 challenge in Syrian hamsters.

At least four different virally vectored vaccines have been tested in Syrian hamsters [20, 21]. A single dose of IN delivered ChAd-SARS-CoV-2-S vaccine induced serum antiviral neutralizing titers of around 1:1217, which is at least several fold higher than IM delivery of Ad26-S (1:360) and VSV-S, or intraperitoneal delivery of Y17F-S (1:630). Furthermore, IN delivery of ChAd-SARS-CoV-2-S protected the upper respiratory tract against infection with SARS-CoV-2 and no weight loss was detected after virus challenge. This contrasts with the other vaccine candidates where the vaccinated hamsters lose between 0 and 5% of their body weight after challenge.

The reason for the higher antibody responses after IN versus IM immunization currently is not known. One possibility is that the respiratory tract of the hamster is more permissive for the ChAd virus than muscle, which could increase the amount of SARS-CoV-2-S antigen produced. Alternatively, the mucosal immune response to the S protein or ChAd infection in the respiratory tract is unique compared to the thigh muscle. The most striking difference between IN and IM immunization is the enhanced protection of the upper respiratory tract infection, with minimal or no viral RNA detected in the nasal olfactory neuroepithelium, which expresses known receptors for SARS-CoV-2 [26]. This effect may be due to the induction of local S protein specific immunity capable of neutralizing virus in the upper respiratory tract. In mice, IN immunization induced robust S-specific IgA antibodies [23], whereas IM immunization did not. Anti-hamster-IgA secondary antibodies currently are not commercially available. Nonetheless, we would expect to find similar IgA responses that can neutralize incoming virus. As a result of the superior protection of the nasal cavity and upper respiratory tract, it IN delivery of ChAd-SARS-CoV-2-S may offer protection against both infection and transmission of SARS-CoV-2.

IN immunization offers many benefits over more traditional approaches [27]. Besides the ease of administration and lack of needles, IN delivery is associated with mucosal immune responses including the production of IgA and stimulation of T- and B-cells in the nasopharynx-associated lymphoid tissue [28]. Influenza virus vaccines are the only licensed IN vaccines to date for individuals over the age of 2 and less than 50 years old. Live-attenuated influenza virus vaccine (LAIV) are considered safe and efficacious. The exception to this was the 2013-2014 and 2015-2016 season when the vaccine was no effective against one of the four components [29]. IN delivery of several viral vectored vaccines has been evaluated in pre-clinical animal models. A single dose of chimpanzee adenovirus vectored vaccine against the Middle Eastern Respiratory Syndrome virus protected hDPP4 knock-in mice and rhesus macaques from MERS challenge [30, 31]. Similarly a parainfluenza virus 5 vectored vaccine expressing the S protein of MERS protected mice from MERS [32]. A replication-incompetent recombinant serotype 5 adenovirus, Ad5-S-nb2, carrying a codon-optimized gene encoding Spike protein (S), protected rhesus macaques from SARS-CoV-2 challenge [33]. Besides coronaviruses, IN delivered adenoviral vectored vaccines protected non-human primates from Ebola virus [34]. Importantly, in that study, protection occurred in the presence existing adenovirus-specific immunity. Besides the many advantages of IN vaccines, IN delivery of a replication defective adenovirus 5 vectored vaccines caused infection of olfactory nerves in mice [35]. In humans, IN delivery of a non-replicating adenovirus-vectored influenza vaccine was well tolerated and immunogenic [36].

The pathogenesis following SARS-CoV-2 infection is mediated in part by a pathological inflammatory immune response [1]. To evaluate the efficacy of this vaccine on reducing this part of the syndrome, we quantified changes in gene expression of 22 different hamster inflammatory and immune genes. Eight out of the 22 showed demonstrated a >2-fold increase in gene expression, with a clear enrichment for type I and III IFN-induced genes. The expression of several other hamster host genes, including IFN-γ, interleukins (IL-10 is an exception), TNF-α, and complement factors did not increase after infection. This lack of expression may be due to the time of organ collection (3 dpi), when the inflammatory response is still developing. The lack of IFN-γ could be explained by the increase in IL-10 expression or the early time point evaluated that precedes influx of NK cells and activated T cells.

Correlates of immune protection and SARS-CoV-2 associated disease were investigated in our cohort of hamsters. Virus neutralization in serum correlated better with RBD-specific antibody levels (r = 0.83, P < 0.0001) than S-protein specific responses (r = 0.31, *P* > 0.05, **Fig S6**). Of the three humoral response parameters, the virus neutralization serum titer (FRNT, IC_50_) correlated best with weight loss 3 dpi (r = 0.59, *P* < 0.0001, **Fig S6**). Weight loss at 3 dpi was strongly associated with viral RNA levels (r = −0.68, *P* < 0.001) and infectious virus load (r = −0.62, *P* < 0.01, **Fig S6**) in the lungs, but not in the nasal swabs. Finally, inflammatory host gene-expression (*Ifit3* and *Cxcl10*) correlated with RNA levels in the lungs (r = 0.68 and 0.84 respectively, *P* < 0.001), and serum virus neutralization titers (r = −0.70 and −0.64 respectively, *P* < 0.01). These analyses suggests that RBD-specific, but not S-specific, serum antibody and virus neutralization titers are important parameters of protection against SARS-CoV-2 and COVID-19 and that high antibody levels are associated with protection from infection and inflammation.

Overall, our studies in hamsters demonstrate that IN delivery of the ChAd-SARS-CoV-2-S vaccine confers protection against SARS-CoV-2 challenge. Protection is associated with lower virus levels in the lungs and upper and lower respiratory tracts, no weight loss, and reduced inflammation in the lungs. These findings support further pre-clinical and clinical studies investigating the vaccine efficacy of IN delivered vaccines against SARS-CoV-2.

## ACKNOWLEDGEMENTS

We acknowledge Krzysztof Hyrc for help with the Hamamatsu NanoZoomer slide scanning system. We also thank Amy Dillard for help with the serum collections from the animals and Ali Ellebedy and Jackson Turner for help with antibody ELISA and recombinant protein. We thank Drs. Susan Cook and Ken Boschert for support of the COVID-19 research. This study was funded by NIH contracts and grants (R01 AI157155, 75N93019C00062, U01 AI151810, HHSN272201400018C, and HHSN272201700060C). J.B.C. is supported by a Helen Hay Whitney Foundation postdoctoral fellowship.

## AUTHOR CONTRIBUTIONS

T.L.B., and T.L.D. performed the animal experiments. A.O.H. upscaled and purified the ChAd vectors. A.O.H., A.C.M.B., and J.B.C. performed the serological analysis. H.H., T.L.B., and T.L.D. performed viral load analysis by quantitative RT-PCR, focus forming or plaque assay. T.L.D. developed and validated primer-probes sets for Syrian hamster genes and performed the host gene expression assays. M. J. H. performed the histology analysis. A.S., X.J. and R.K. performed the histology and RNA-ISH on the hamster heads. A.C.M.B. performed RNA-ISH on the hamster lungs. A.L.B. and J.B.C. provided quantitative PCR reagents and protocols. Y.N.D., H.Z., L.J.A., and D.H.F. provided recombinant proteins for serological analysis. M.S.D. and A.C.M.B. provided supervision and acquired funding. A.C.M.B. wrote the initial draft, with the other authors providing editorial comments.

## SUPPLEMENTARY FIGURES

**Supplementary Figure 1:**
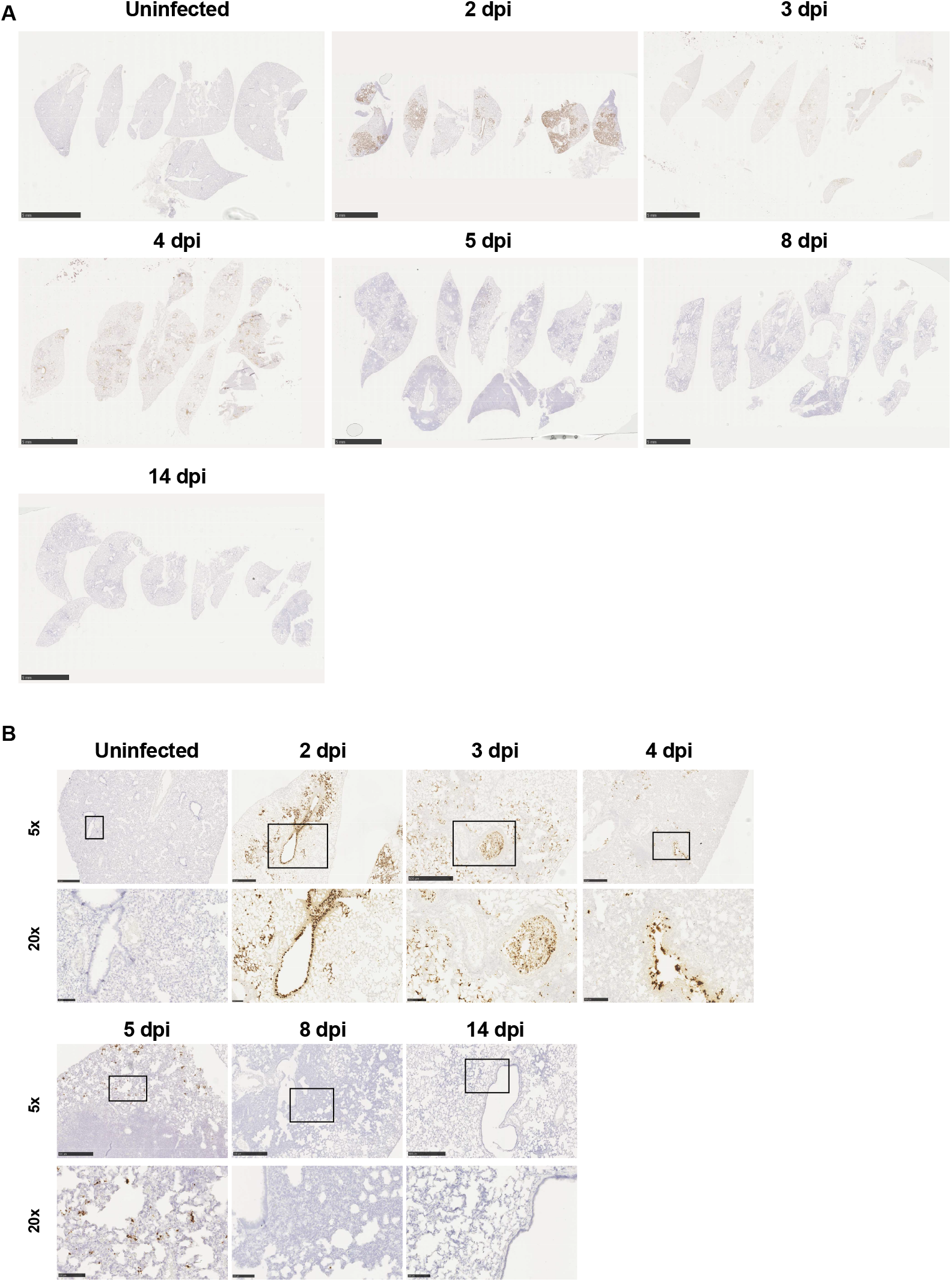
RNA *in situ* (ISH) hybridization on lung tissue sections from SARS-CoV-2 infected hamsters. Representative images at 0.5x (**A**), 5x (**B**), and 20x (**B**) magnification of RNA-ISH of SARS-CoV-2 infected hamsters sacrificed at different time points after inoculation (n = 5 for 2 dpi, n = 3 for 3 dpi, n = 3 for 4 dpi, n = 5 for 5 dpi, n = 2 for 8 dpi, n = 3 for 14 dpi, n = 3 for uninfected).

**Supplementary Figure 2:**
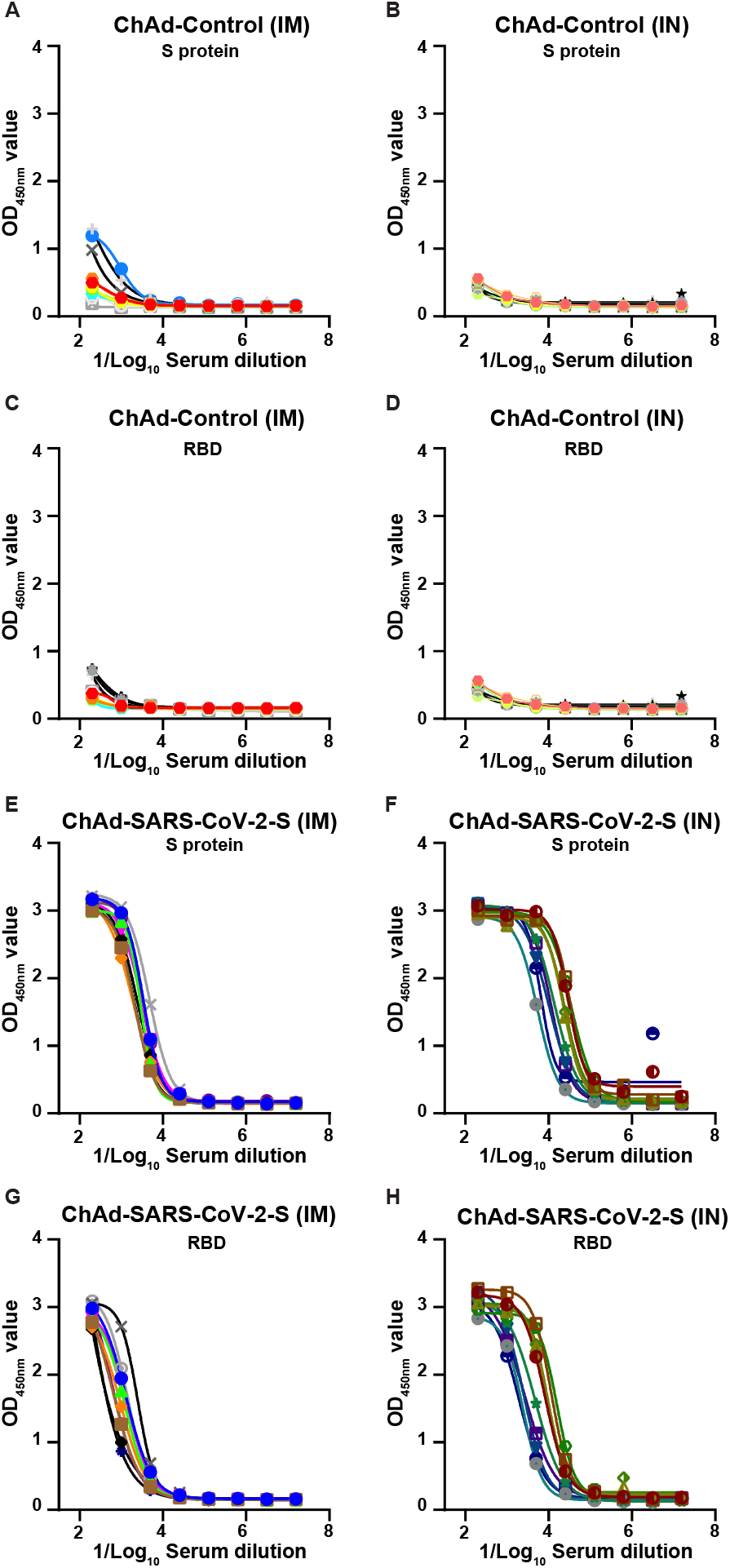
IgG2/IgG3 serum antibody titers against recombinant spike protein and the receptor binding domain (RBD) of the spike protein. (**A-B**) S protein-specific serum IgG2/IgG3 titers in hamsters immunized IM (**A**) or IN (**B**) with ChAd-Control. (**C-D**) RBD-specific serum IgG2/IgG3 titers in hamsters immunized IM (**C**) or IN (**D**) with ChAd-Control. (**E-F**) S protein-specific serum IgG2/IgG3 titers in hamsters immunized IM (**E**) or IN (**F**) with ChAd-SARS-CoV-2-S. (**G-H**) RBD-specific serum IgG2/IgG3 titers in hamsters immunized IM (**C**) or IN (**D**) with ChAd-SARS-CoV-2-S. Each line is an individual animal.

**Supplementary Figure 3:**
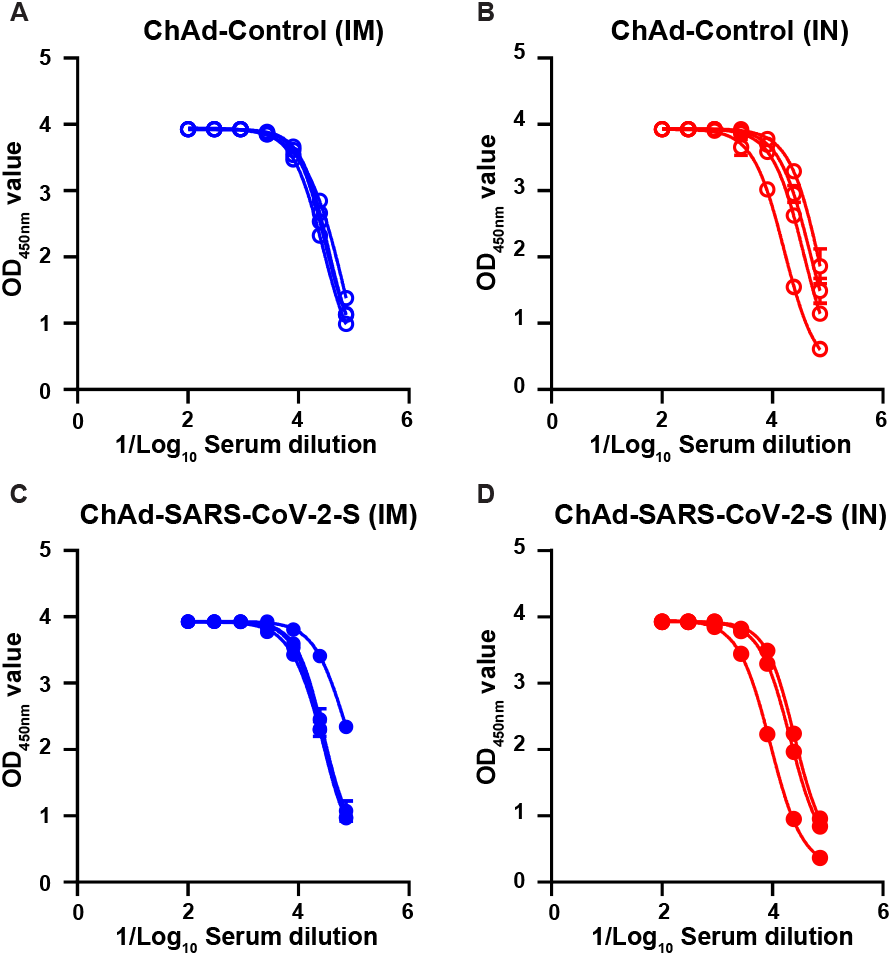
IgG(H+L) serum antibody titers against SARS-CoV-2 nucleoprotein 10 days after infection in vaccinated and control hamsters. Nucleoprotein-specific serum antibody titers in ChAd-Control (**A-B**) or ChAd-SARS-CoV-2-S (**C-D**) immunized and SARS-CoV-2 challenged Golden Syrian hamsters 10 days post challenge. Each line is an individual animal.

**Supplementary Figure 4:**
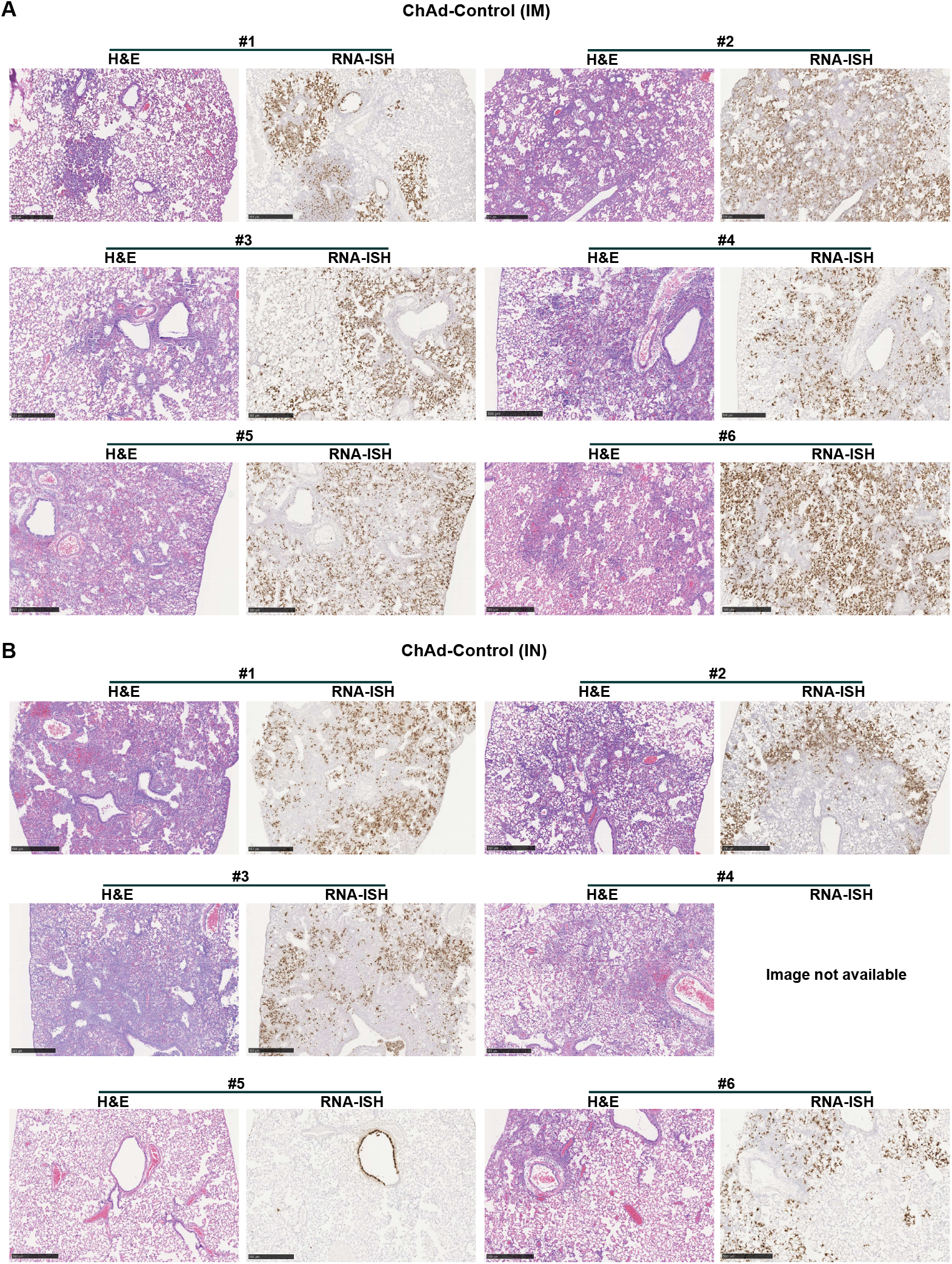
Histological and RNA *in situ* (ISH) hybridization analysis of lung tissue sections from ChAd-Control vaccinated and SARS-CoV-2 challenge hamsters. Representative images at 5x magnification of H & E staining and RNA-ISH of hamsters immunized IM (n = 6) and IN (n = 6) with ChAd-Control and challenged 28 days later with SARS-CoV-2. Lungs were collected 3 days post challenge, fixed in 10% formalin and paraffin embedded prior to sectioning and staining.

**Supplementary Figure 5:**
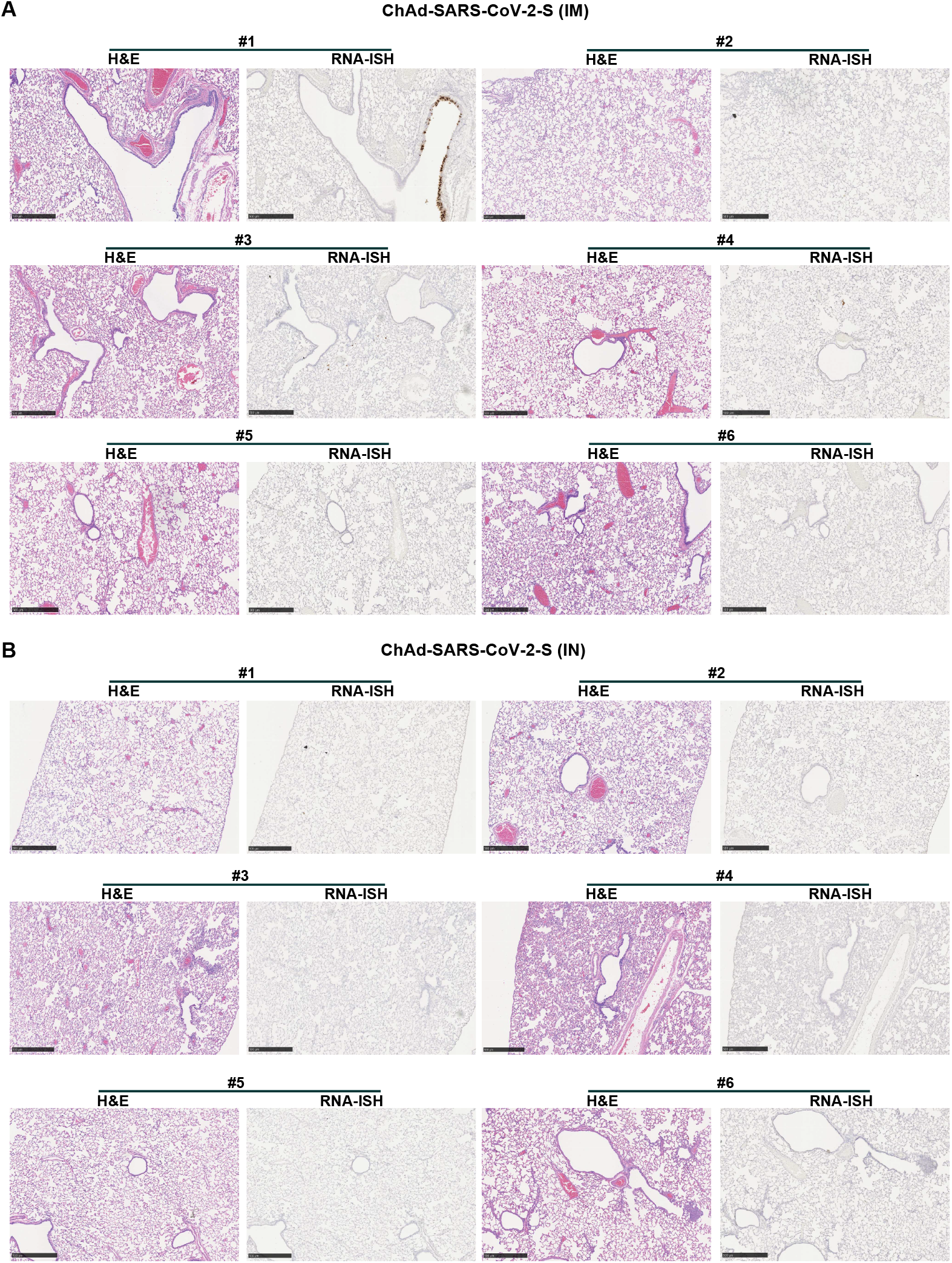
Histological and RNA *in situ* (ISH) hybridization analysis of lung tissue sections from ChAd-SARS-CoV-2-S vaccinated and SARS-CoV-2 challenge hamsters. Representative images at 5x magnification of H & E staining and RNA-ISH of hamsters immunized IM (n = 6) and IN (n = 6) with ChAd-SARS-CoV-2S and challenged 28 days later with SARS-CoV-2. Lungs were collected 3 days post challenge, fixed in 10% formalin and paraffin embedded prior to sectioning and staining.

**Supplementary Figure 6:**
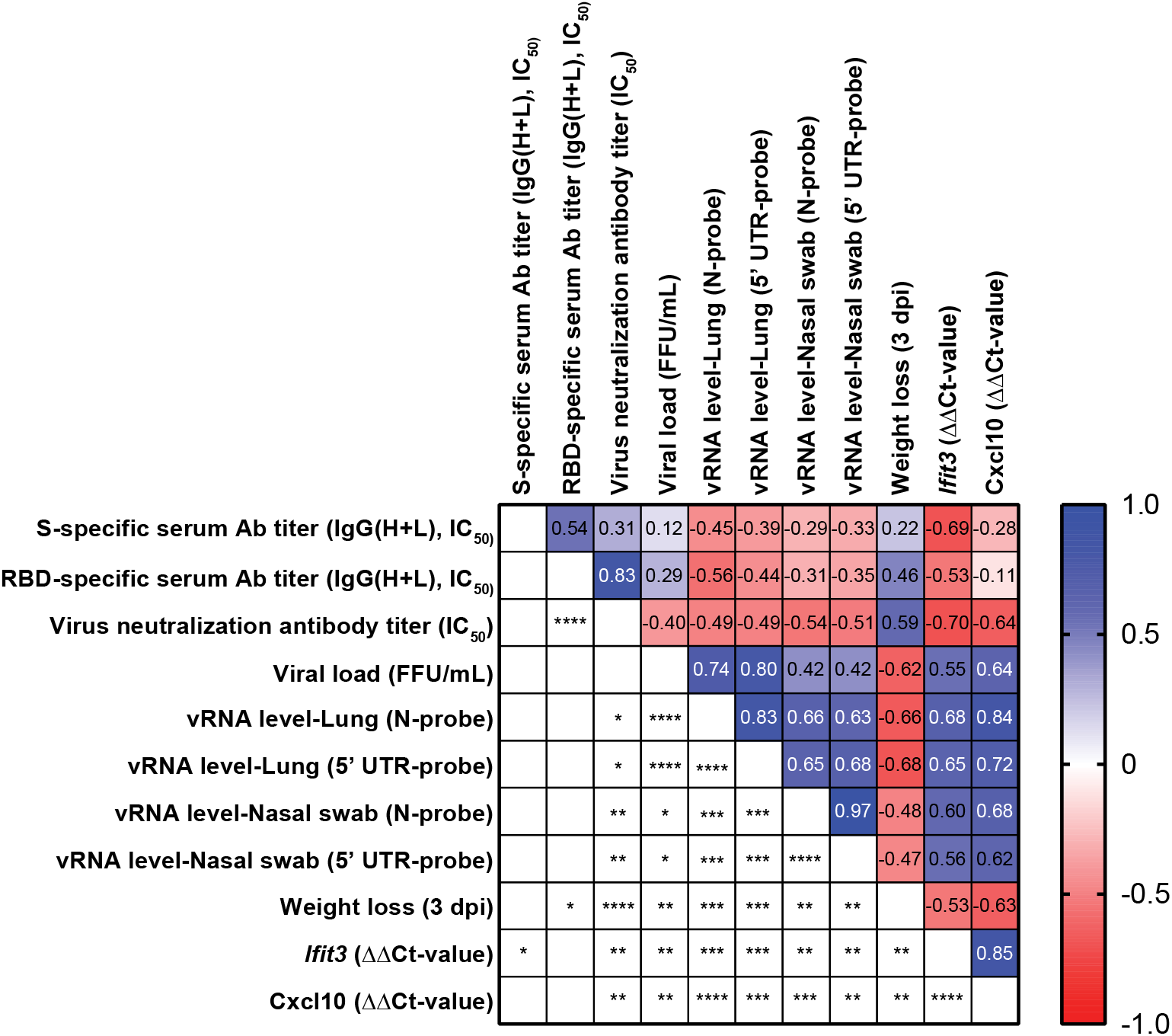
Immune correlates of vaccine-mediated protection SARS-CoV-2. Correlations between % weight-loss/gain (3 dpi), RNA levels in the lungs and nasal swabs, infectious virus titers, serum antibody responses, serum virus neutralization titer, and inflammatory hamster gene expression were analyzed for all animals in the vaccine study using a Pearson correlation matrix. The top right side is the correlation coefficient and the bottom left side has the P-value for every combination (**** *P* < 0.0001, *** *P* < 0.001, ** *P* < 0.01, * *P* < 0.05 by Pearson’s correlation analysis).

## SUPPLEMENTARY TABLES

**Supplementary Table 1:**
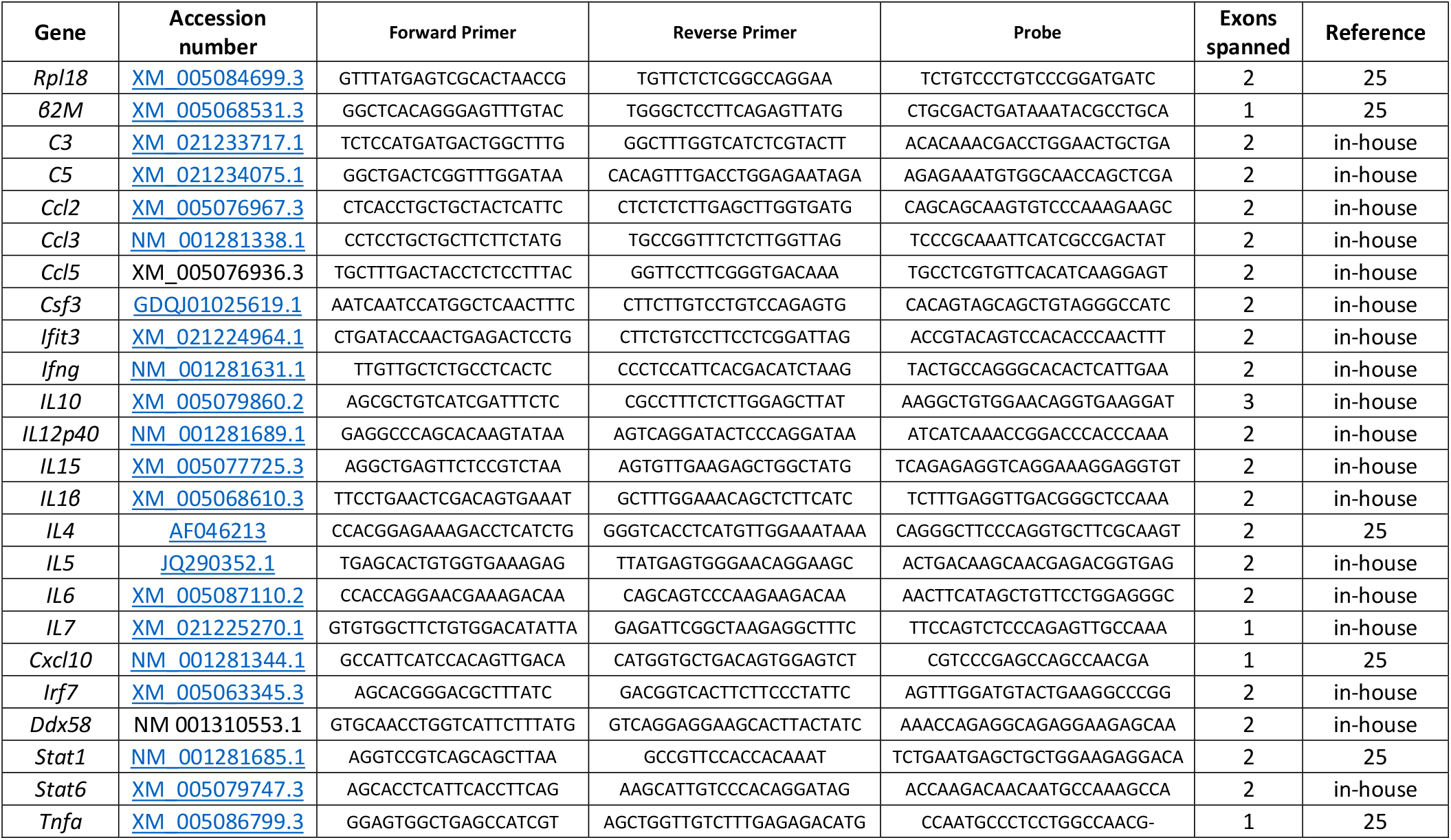
Primers and probe sets used to quantify gene expression in the Golden Syrian hamster (*Mesocricetus auratus*).

